# Can Pupillometry Reveal Perturbation Detection in Sensorimotor Adaptation during Grasping?

**DOI:** 10.1101/2025.05.15.654224

**Authors:** Luise Pfalz, Carl Müller, Karl Kopiske

**Author notes:** Corresponding author: Karl Kopiske.

## Abstract

Humans adjust their motor actions to correct for errors both with and without being aware of doing so. Little is known, however, about what makes errors detectable for the actor. Here, we replicate and extend prior work showing that motor adjustments may mask the very errors they correct for. We also investigated pupillometry as an unobtrusive no-report marker of perturbation detection. N=48 participants grasped objects while a visuo-haptic size mismatch was applied either sinusoidally or abruptly. When mismatches started abruptly and thereafter stayed the same, participants adapted well but also showed decreasing discrimination performance and decreasing confidence in their responses. This was not the case for sinusoidally introduced perturbations. We also show that parameters that characterize phasic and tonic pupil responses were predicted by stimulus parameters and differed depending on participants’ grasping and behavioral responses. However, predicting response characteristics from pupil-dilation features using support-vector machine classifiers was not successful. This shows that while pupillometry may yet prove to be a useful no-report marker of perturbation and error detection, there are some challenges for trial-by-trial prediction.

**New and Noteworthy:** An actor detecting the need to adjust a motor action can make these adjustments more efficient. We show that error signals play a central role in both humans’ detection of and meta-cognition about motor perturbations. Pupil dilation during perturbed actions reflected perturbation properties and participants’ responses, but trial-wise prediction of responses using pupil-dilation parameters was close to chance. This is a step towards determining whether pupillometry can serve as a no-report marker of perturbation detection.

## Introduction

### Visuomotor mapping and forward mechanisms in motor actions

Imagine that a friend brings you an unusually shaped, extravagant cup from their vacation (Fig. 1). The handle is positioned slightly upper, the diameter is larger, and the shape differs significantly from your usual cup, shifting its center of mass. As you drink your morning coffee, you notice that you frequently misreach, fail to grasp the handle correctly, or bump your hand against the rim. It becomes clear: you need to adapt to this new cup. But why does this process seem so challenging?

**Figure 1:**
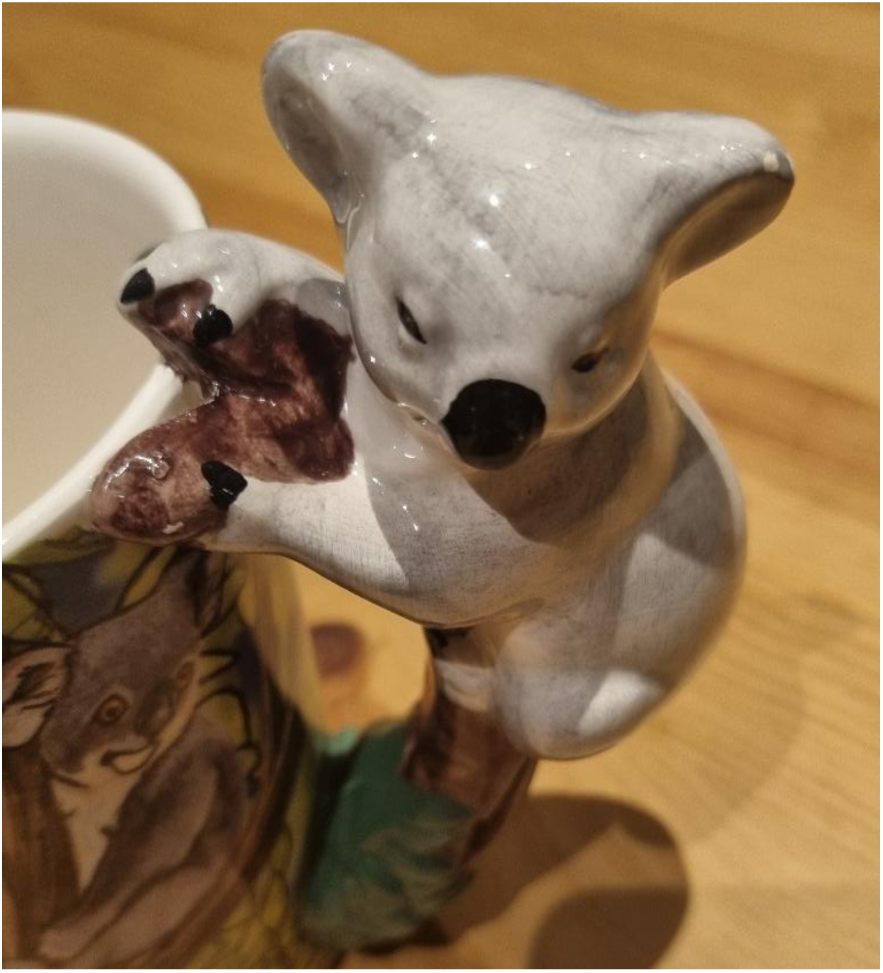
An unusual coffee cup.

The key to this improvement is sensorimotor adaptation. This process describes how the brain adjusts movements such as grasping actions to altered conditions (Krakauer & Mazzoni, 2011). Several components must interact on a physical level: joints in the shoulder, elbow, wrist, and fingers need to be precisely coordinated to securely grasp the cup. Simultaneously, the physical properties of the cup – its mass, shape, orientation, and size – affect hand positioning. This physical adjustment is guided by a neural system that translates perceptual information into motor commands, a process known as perceptual-motor mapping (Warren, 2006).

Computationally, it is typically modelled with so-called internal forward mechanisms: Based on the current position and movement of the hand, the brain generates a prediction of where the hand will land on the object – in this case, on the handle of the new cup. This prediction is compared with sensory feedback, which provides information on where the hand actually landed on the object (Miall & Wolpert, 1996). If there is a mismatch, the brain receives an error signal, and an adjustment is made to avoid the error in the future. This explains why grasping the new coffee cup becomes easier with each attempt: The mapping is updated with each movement, improving the grasp over time and increasing the likelihood of a successful, fluid grasp (Wolpert et al., 1995). Eventually, your movements become as accurate as before.

### Implicit and explicit processes in adaptation

The adjustment or correction of the grasping movement is considered in large part as an implicit learning process, that is, outside of cognitive control (Mazzoni & Krakauer, 2006) and controlled largely by forward models in the cerebellum (Shadmehr et al., 2010). However, explicit, deliberately controlled processes have also been shown to influence motor adaptation (Taylor & Ivry, 2011), allowing for rapid, consciously controlled or even strategic adjustments to correct for errors, with central involvement of memory areas such as the medial temporal lobe (McDougle et al., 2022). In contrast, implicit learning proceeds more slowly, and without the actor being consciously aware of them. These processes work in tandem, responding to different error signals – both the conscious correction after the first attempt and the unconscious adaptation that occurs over time – and together they optimize motor adaptation to perturbations (McDougle et al., 2016; Taylor et al., 2014).

In experimental research, these two processes have been separated by specific experimental manipulations, such as distracting participants from the presence of perturbations (Mariscal et al., 2020) or by providing them with explicit instructions to compensate for perturbations (Miyamoto et al., 2020; Taylor & Ivry, 2011). However, in real-world situations, no instructions are provided, and motor errors are detected based only on their inherent characteristics – if at all. Research has investigated adaptive behaviors when perturbations were intentionally made very large (Hudson & Landy, 2012), when participants received identical feedback regardless of whether their movement was accurate or erroneous, known as a “clamp” perturbation (McDougle et al., 2015), or when perturbations were introduced abruptly rather than gradually (Modchalingam et al., 2023; Orban de Xivry et al., 2013). In all these cases, conditions were chosen such that perturbations should be detected in one condition but not in another (or at least far more frequently in one condition). Much less research, however, has investigated when participants can perceive inherent properties of perturbations.

One example of this is a study by Gaffin-Cahn and colleagues (Gaffin-Cahn et al., 2019). Here, participants performed a reaching task with distorted endpoint feedback and judged whether the endpoint shown to them was the result of their own movement. Participants were able to identify the perturbations, relying primarily on visual endpoint feedback and less on proprioceptive cues, which indicates that different external factors are weighed differently. Other studies have investigated metacognitive judgements about motor errors: That is, the extent to which participants notice and are confident about motor errors during their movements. Pereira and colleagues (Pereira et al., 2023) had participants perform a target task using a joystick, while deviations in cursor movement were experimentally introduced. Participants indicated if they had noticed a deviation and rated their confidence in their decisions while neural activity was recorded using fMRI. Results showed that participants could adjust confidence in their decisions according to the accuracy of their responses, even when they were unaware of the deviations. Similarly, Arbuzova and colleagues (Arbuzova et al., 2021) showed metacognitive awareness of motor errors during a virtual ball-throwing task. This suggests a level of metacognitive awareness based on visuomotor information even in the absence of conscious awareness of errors, although error history might play a crucial role in it (Hewitson et al., 2023).

To examine directly which factors affect perturbation detection, a previous study that our experiments are based on (Müller et al., 2025) investigated how a size mismatch between a visible and an invisible target cuboid, as well as the sensory error signal (the difference between the expected and actual haptic grip feedback), affect motor adaptation and mismatch discrimination in a grasping task. Two types of perturbations were used: an abrupt perturbation, where the mismatch between cuboids was introduced in the first trial and then remained constant, and a sinusoidal perturbation, where the mismatch varied in each trial. Participants were instructed to indicate in a two-alternative forced-choice (2AFC) task whether the grasped cuboid was larger or smaller than the visible one, founding that discrimination performance gradually declined when participants adapted their grip to the mismatch for abrupt perturbations. However, with a continuously changing mismatch (sinusoidal) without systematically decreasing error signal, discrimination performance remained constant.

### Pupillometry as a physiological measure of perception

To directly capture participants’ perception of a perturbation, they must be asked (e.g., “Was the object you saw smaller or larger than the one you felt?”, or “Where will your pointing movement land?”). However, this can be challenging, as those same questions alert participants to the perturbations, and with typical adaptation schedules, repeated responses might cause uncertainty, influencing the participants’ answers (Bosch et al., 2020). Therefore, it may be useful to employ alternative methods espousing direct responses from the participant, such as physiological correlates.

Pupillometry has served as a way around requiring participants to report their perception altogether and to circumvent the issues that come with this. For example, Einhäuser and colleagues (Einhäuser et al., 2008) demonstrated that pupil responses could serve as an indicator of perceptual selection in multistability, such as the perception of a Necker cube, as pupil size correlated with perceptual changes. It has also been used as a no-report marker of perception in binocular rivalry (Naber et al., 2011). Results like these suggest that pupil size may reflect changes in visual perception and cognitive processes. Similarly, Yokoi and Weiler (Yokoi & Weiler, 2022) examined changes in pupil size during motor adaptation. Participants performed reaching movements using a robotic manipulandum, where a lateral force was introduced as a perturbation through a force-field perturbation. These perturbations were applied either abruptly with multiple direction changes, or gradually. To assess the effects on pupil dilation of uncertainty and surprise, the authors focused on trials following the introduction of the perturbation, and after the perturbation was removed, but without validating the participants’ subjective perception. The results indicated that phasic pupil response, that is, the baseline pupil diameter, was significantly increased at the onset of each experimental block, also associated with prolonged reaction and movement times. A pronounced tonic pupillary response, that is, change in dilation during the movement, was observed both at the start of a new block and during the introduction of a perturbation. The authors concluded that not only physical exertion but also the perturbation itself triggers a pupillary response perhaps related to locus coeruleus activity in response to surprise (Dayan & Yu, 2006; Yokoi & Weiler, 2022). This supports the notion that larger pupil diameters and faster pupil reactions are associated with increased uncertainty and more difficult perceptual decisions and that pupil size correlates with subjective confidence and surprise about environmental changes.

In summary, participants can detect perturbations during grasping tasks, and their pupils respond to perturbations. This raises the question of whether pupillary responses alone could be used to predict the participants’ current perception – without requiring explicit feedback from the participants. The aim of this study was to tackle this question by combining (a) a grasping task with different perturbation schedules with (b) a psychophysics task where participants judged the relative sizes of visual and haptic (i.e., seen and felt) objects and (c) pupil dilation during the tasks, which was evaluated depending on stimulus and response characteristics and used to predict the response.

## Methods

### Participants

A total of 59 participants took part in this study. Participants were required to be between 18 and 60 years old, right-handed, and have unrestricted arm and hand functionality as well as normal or corrected-to-normal vision, provided any visual aids needed did not interfere with the participant’s comfort or the measurement process. Of the 59 participants, 11 were excluded from the experiment due to technical difficulties with achieving a successful eye-tracker calibration through the PLATO goggles and the cold mirror, resulting in 48 participants included in the final analysis, the intended sample size to enable counterbalancing of conditions and provide at least 90% power with an effect of Cohen’s d = 0.5 (Cohen, 1988). Of these participants, 33 were women and 15 men, with ages ranging from 18 to 37 years. Participants were compensated either with course credit or 10€ per hour.

### Materials and stimuli

Participants were presented with cuboids of varying sizes in a mirror setup (Müller et al., 2025), see Fig. 2. As was done in Müller et al. (2025), three cuboids were positioned on a rotating platform that could rotate one of them to face the participant on any given trial. The aluminum cuboids used in this experiment had a base of 15x15 and different lengths, with the smallest cuboid measuring 28 mm and the largest 60 mm. The three visually presented stimuli had lengths of 40 mm, 44 mm, or 48 mm. The size of the haptically presented stimuli varied based on the visually presented size and the visuo-haptic mismatch determined by the perturbation condition. This mismatch was introduced either abruptly or sinusoidally, with six different perturbation magnitudes: −12 mm, −6 mm, −3 mm, 3 mm, 6 mm, and 12 mm. Visual and haptic size was dissociated by projecting the “visual” cuboid towards the participant via a cold mirror slanted 45° away from them, while a “haptic” cuboid to be grasped was placed behind the mirror at the location where the participant saw the visual cuboid.

**Figure 2:**
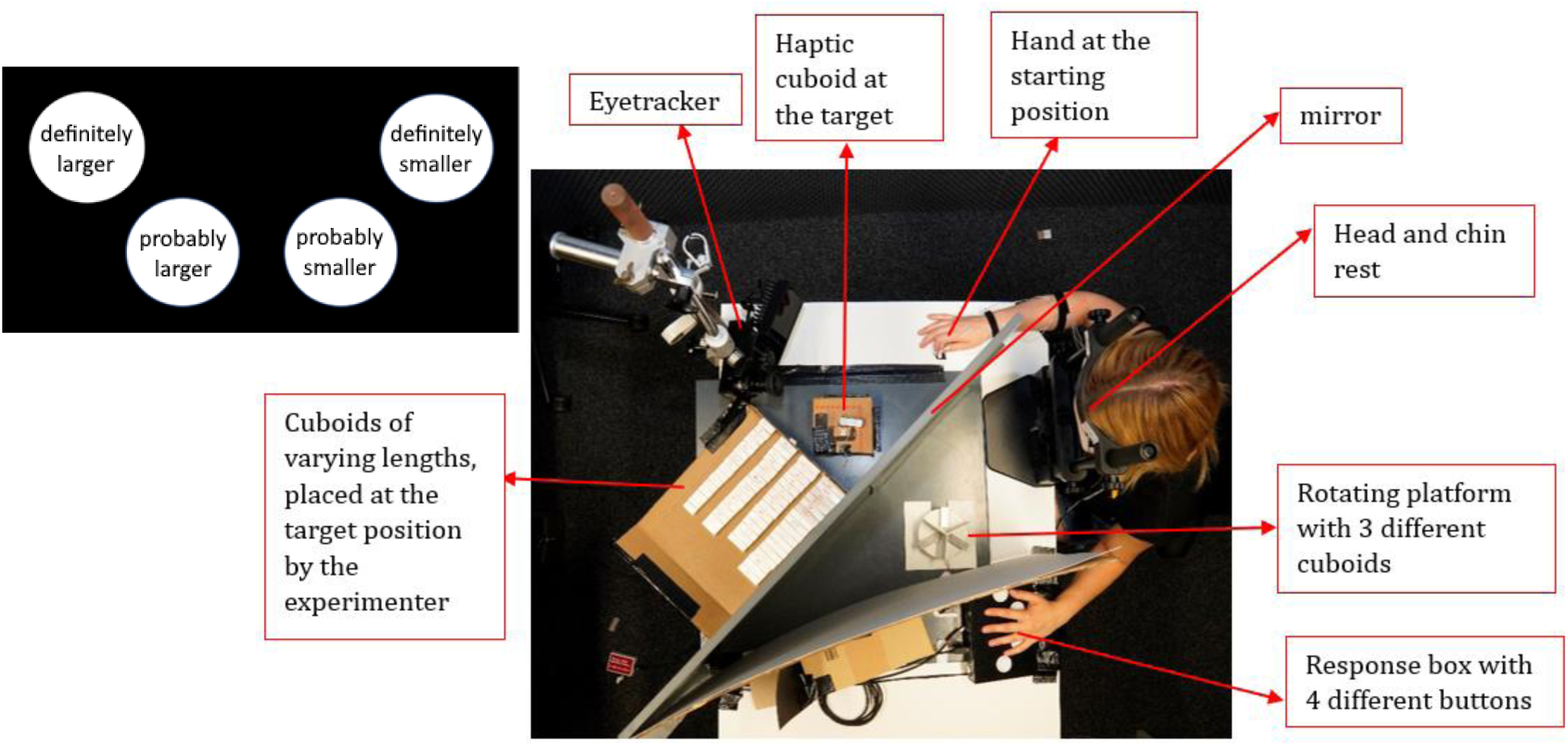
The experimental setup. Participants sat in front of a cold mirror, slanted away from them at 45°, with their head in a chin rest and their left hand on a response box. Visual objects were presented on a rotating platform in front of the mirror, haptic objects were placed behind the mirror in the same position where participants saw the visual objects. PLATO goggles were used to prevent vision of the platform rotating, an EyeLink-1000 tracked gaze position and pupil diameter through the cold mirror. Inset, top left: The responses assigned to the four buttons of the response box.

Participants wore PLATO goggles (Milgram, 1987) to remove vision between trials, whose opaque lenses obstructed vision when closed but minimally altered ambient brightness (to 80-90%), thereby only slightly affecting pupil dilation. Differing from Müller et al. (2025), responses were given using a four-button response box (“Black Box”; The Black Box ToolKit Ltd., Sheffield, UK; Fig, 2, inset). Behind the mirror was an EyeLink-1000 eye tracker (SR Research, Ottawa, Canada) tracking participants’ eye movements and pupil dilation through the cold (infrared-transparent) mirror, and an Optotrak 3D Investigator (Northern Digital, Waterloo, CA) that captured hand movements at a frequency of 500 Hz. Infrared diodes were placed on the participants’ thumb, index finger, and wrist. To determine the exact moment of contact with the object, reflective aluminum was affixed to the long sides of the haptic object, and a diode was positioned nearby to allow the diode’s signal to be reflected. Once the haptic cuboid was lifted, the signal was no longer reflected.

### Experimental design

For each trial, participants were instructed to grasp the haptic cuboid, lift it briefly, place it back down, and then judge whether the haptic cuboid was larger or smaller than the visual cuboid. Four response options were provided: “definitely larger,” “probably larger,” “probably smaller,” and “definitely smaller”, allowing more nuanced insights compared to a simple “larger or smaller” decision (Müller et al., 2025) and including subjective confidence as a factor in pupil-dilation analyses – that is, we considered “definitely larger” and “definitely smaller” as responses with high subjective confidence, and “probably larger” and “probably smaller” as low-subjective confidence responses. This is in line with previous work showing uncertainty about the perturbation on the next action being a predictor of pupillary responses (Yokoi & Weiler, 2022). We deliberately included confidence in one single judgement on each trial despite concerns that this can introduce biases (Fleming & Lau, 2014; Mamassian, 2016), since this type of response ameliorated a key issue in Müller et al. (2025), participants repeatedly having to give the same response to repeated identical perturbations.

Prior to the main experiment, a training block consisting of 12 trials was conducted to familiarize participants with the task. This training block contained both unperturbed trials and trials with the largest positive and negative size mismatch. During training, participants were informed when they had grasped a perturbed object, helping them develop an understanding of the potential size differences. In the main experiment, the abrupt-perturbation condition was presented a total of 6 times (±3 mm, ±6 mm, ±12 mm), and the sinusoidal-perturbation condition was presented 3 times (3 mm, 6 mm, 12 mm). To control for sequential effects across blocks, a 6x6 row-balanced Latin Square was used to determine the order of the abrupt blocks, while the sinusoidal blocks were inserted at randomly selected positions. The order of these blocks also varied across participants. Consequently, each participant completed 9 blocks, each lasting approximately 11 minutes (abrupt blocks ∼10 minutes, sinusoidal blocks ∼12 minutes). After each block, participants were allowed to take a break for as long as needed.

Each block in the *abrupt-perturbation* condition consisted of 24 trials. The first four trials presented blocks identical to the visually displayed ones to establish a baseline for the maximum grip aperture (MGA). In the subsequent 16 trials, the length of the haptic cuboid was either larger or smaller than that of the visual cuboid by a fixed amount, depending on the perturbation magnitude assigned to the block. The rotating platform displayed a different cuboid in each trial, with the size mismatch being constant across each one block of the abrupt-perturbation condition. For instance, if the visual cuboid in a perturbed trial had a size of 44 mm and the perturbation magnitude was 6 mm, the haptic cuboid had a size of 50 mm. The final four trials were washout trials in which the perturbation was removed, and the visual and haptic cuboid were of equal size again. This allowed for the assessment of the aftereffects of the perturbation on the MGA and the psychophysical judgments.

Each block in the *sinusoidal-perturbation* condition consisted of 36 trials. The length of the haptic cuboid was altered according to a sinusoidal function (as proposed by Hudson & Landy, 2012) with each sinus-cycle consisting of 12 trials, thus creating size differences between the visual and haptic objects without abrupt changes. As a previous study (Müller et al., 2025) found no significant difference in response patterns between positive and negative perturbation magnitudes, these were combined in the current study, and each participant completed three sinusoidal-perturbation blocks with maximal amplitudes of 3 mm, 6 mm, and 12 mm, respectively. A randomized phase shift was introduced, starting the sinusoidal cycle in either the positive or negative direction. Across all participants, an equal number of positive and negative perturbation amplitudes were presented, and each participant completed at least one of each.

### Data processing and analysis

#### Grasping: Maximum grip aperture and adaptation

To filter the motion-tracking data, we applied a cubic-spline interpolation and a third-order Savitzky-Golay filter (Savitzky & Golay, 1964) using a 55-ms window. For each trial, the onset of the grasping movement was defined as both the index finger’s and thumb’s marker moving at more than 25 cm/s, while the end was defined as the moment when the diode that was reflected by the target object was no longer visible to the motion-capture system or moved at more than 25 cm/s. Trials were excluded from analysis if the grip-aperture trajectory was missing more than 20% of frames, or as outliers if the MGA was more than three inter-quartile ranges removed from a participant’s median for the respective visual object size (3.8% of trials combined).

To measure grip adjustment to the different cuboid sizes, the distance between the thumb and index finger was recorded and their maximal distance during the grasping movement was computed as the MGA. The MGA is known to increase with object size (Bhatia et al., 2022; Jeannerod, 1984; Smeets & Brenner, 1999) and reflects sensorimotor adaptation when participants adjust their grip to visual or haptic perturbations (Cesanek & Domini, 2017; Gentilucci et al., 1995; Kopiske et al., 2017; Säfström & Edin, 2005). We modelled adaptation of the MGA in response to the error signal (the size difference between the visual and haptic cuboids) using a linear state-space model (Cheng & Sabes, 2006; Wolpert et al., 1995) in which a state, corresponding roughly to a visuomotor mapping of visual input to a motor action, changes linearly from the previous state based on the error signal from the previous trial, thereby facilitating a correction or adaptation in the grip movement:

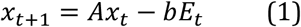

Here 𝑥_𝑡_ represents the current state and is modified trial-by-trial based on the error 𝐸_𝑡_, which we defined as the haptic error signal – that is, difference between the observed MGA and the MGA predicted from the linear response function of MGA ∼ haptic size and the haptic object size on each trial. The retention parameter A indicates the extent to which the previous state influences the current state. For fitting, the nloptr package (Ypma, 2014) was used. Parameters A and b were each bounded between [0, 1].

To assess the adaptation process, the correction parameter b was our main parameter of interest. To assess whether adaptation differed between conditions, we conducted a repeated-measures ANOVA (rmANOVA) with factors *perturbation schedule* (abrupt or sinusoidal) and *perturbation magnitude* (3 mm, 6 mm, 12 mm).

#### Psychophysics: Size discrimination

We analyzed perturbation-detection, measured indirectly through size discrimination, in three ways: One, overall performance was assessed by creating receiver-operator-characteristics (ROC) curves (Green & Swets, 1966) for each participant and each perturbation schedule, with each of the four response levels to the question “was the felt object larger or smaller than the seen one?” essentially being treated as different decision criteria (Naber et al., 2013). That is, each curve consisted of four points with y_1_ equaling the proportion of “definitely larger” responses when the haptic object was indeed larger, y_2_ being the combined proportion of “definitely larger” and “probably larger” responses, etc., and the x-coordinates being the corresponding values for smaller haptic objects and responses starting with “definitely smaller”. In sinusoidal blocks, only trials with maximum amplitude were used in this analysis, to enable a fair comparison to abrupt blocks. Two, we collapsed “definitely” and “probably” correct responses and “definitely” and “probably” incorrect responses, respectively, to get a binary correct/incorrect scoring that could be used to (i) replicate the finding from previous work (Müller et al., 2025) and (ii) compute linear slopes for the percentage of correct responses over trials in each block, which served as a means to estimate if participants got better or worse as perturbations were presented repeatedly. These slopes were then submitted to rmANOVAs with factors perturbation schedule (abrupt or sinusoidal) and perturbation magnitude (3mm, 6mm, 12 mm). Three, we conducted these same rmANOVAs for the slopes of response confidence after dividing responses in confident vs. unconfident.

#### Pupillometry: Preprocessing, parameters, prediction

Throughout the experiment, eye movements and pupillary responses were recorded at a frequency of 1000 Hz. Pupil-dilation trajectories from 1000 ms before and 2500 ms after contact with the haptic cuboid were used. Blinks were automatically identified by EyeLink DataView (SR Research, Ottawa, Canada), and all data 50 ms around blinks was removed. Plots and basic analyses were conducted with unfiltered data. To train classifiers, missing data was linearly interpolated and data filtered with a 35-ms Savitzky-Golay filter (Savitzky & Golay, 1964). An average of approximately 42% of data points were missing, including blinks. Visual inspection (Fig. 7) shows that missing data were spread relatively evenly across trajectories and did not cluster around systematically relative to touch, although perhaps somewhat after the response was given (Fig. 7, top row), with mean peaks of 48.6%. Trials were excluded from analysis if less than 500 ms of the trajectory could be evaluated (4.9% of trials). One experimental block in one participant additionally had to be excluded due to technical issues during recording.

From each trial’s pupil-dilation trajectory, we computed a set of three parameters: (i) The baseline (computed as the mean dilation during a 1,000 ms window after opening of the goggles) was computed as a measure of tonic pupil response. (ii) We estimated the dilation’s maximum and minimum velocity (i.e., the maximum speed of the pupil opening and contracting) post-touch and took the difference between the two as a measure of phasic response. (iii) Finally as another measure of phasic response that was also used by (Yokoi & Weiler, 2022), we calculated the amplitude (i.e., peak-trough difference) of the aperture post-touch.

To analyze which variables (trial parameters as well as properties of the grasp and the psychophysical response) influenced the pupillary response, linear mixed-effects models (LMEs) were applied (Bates et al., 2015). We iteratively fit models with an increasing numbers of predictors, discarding factors that did not improve the Akaike Information Criterion, *AIC* (Akaike, 1974; Burnham & Anderson, 2004) relative to the current best model. In order, we performed this procedure with the factors (each fit as a fixed effect with a slope) *perturbation magnitude*, *perturbation type*, *trial number*, *block*, *response accuracy*, and *confidence* in correct responses, and for each of the pupil parameters *baseline* pupil size, post-touch pupil *amplitude*, and post-touch pupil *velocity difference*. A random effect with a random intercept for each participant was also included in every model. For each pupil parameter, the model with the lowest AIC value was ultimately selected as the one that best explained the pupillary response as a physiological measure of perception influenced by the perturbation.

We used a Support Vector Machine (SVM) to determine whether participants’ responses could be inferred from their pupillary responses and other trial information (Boser et al., 1992). While ideally, one would predict both the correctness and confidence of the response, here we focused on the responses’ confidence, given that this was a parameter that participants had direct access to. Further, it has been argued that pupil responses (also in perception-action tasks) are particularly sensitive to surprise (Yokoi & Weiler, 2022), which also might be reflected in the difference between confident and unconfident correct responses. Thus, the following analysis examined the extent to which confidence could be predicted, in correct responses, based on different groups of predictor variables.

We used five sets of features to compute subject-wise SVM classifiers: (i) trial information (block number, trial number, perturbation type, perturbation magnitude), (ii) classic “pupil parameters” as defined above (baseline, velocity difference, amplitude), (iii) pupil-response trajectories spanning from 500 ms before to 2500 ms after contact with the haptic cube, downsampled to 50 Hz (to make computation feasible), (iv) the first temporal derivative of those trajectories, also downsampled to 50 Hz, and (v), the grasping error, defined as the difference between the observed MGA and the MGA expected given the response function and the haptic object size. For each participant, we used seven randomly selected experimental blocks as the training set to train a c-classification SVM with radial-basis kernel using the R-package *e1071* (Meyer et al., 2024), and the remaining two blocks as the test set. Each combination of the four feature classes was used, with the class-balanced accuracy in the test set being our primary outcome. Since classification performance would likely depend to some extent on both the sampling rate of the data and meta parameters of the SVM model (Kuhn & Johnson, 2013), we varied the sampling rate (ranging from 5 Hz to 50 Hz), as well as the cost-parameter of constraints violations (i.e., the factor by which residuals are multiplied, ranging from C = 10^-5^ to C = 10^5^) and the class weights for responses (using both equal weights and weights inverse to the response distribution). We report results from our “standard” combination of parameters with 50Hz, a cost parameter of C = 1, and equal weights for each class of responses in the main text, and the range of model performance based on varying these parameters in the appendix. Data and analyses are available at https://osf.io/vh36g/?view_only=9e72dec48ed04b57a1c24fe5df82df6d

## Results

### Maximum grip aperture and sensorimotor adaptation

Participants scaled their grip to the size of the visual object, mean slope = 0.44 ± a standard error of 0.05. Mean parameters from our error-correction model (eq. 1) were b = .35 ± .01 and A = .80 ± .01. The mean correction parameters by perturbation type were *b_abrupt_* = 0.42 ± .01 for the abrupt condition and *b_sinusoidal_* = 0.21 ± .01 for the sinusoidal condition. This difference was significant in the rmANOVA (*F*(1, 47) = 95.40, *p* < .001), indicating that adaptation to the perturbation was significantly more effective in the abrupt condition than in the sinusoidal condition. There was also a main effect for perturbation magnitudes (*F*(2, 94) = 4.22, *p* = .02), but no interaction (*F*(2, 94) = 3.14, *p* = .097). Grasping data aggregated by perturbation type and magnitude, as well as from two blocks of a sample participant to illustrate the models, can be seen in Fig. 3.

**Figure 3:**
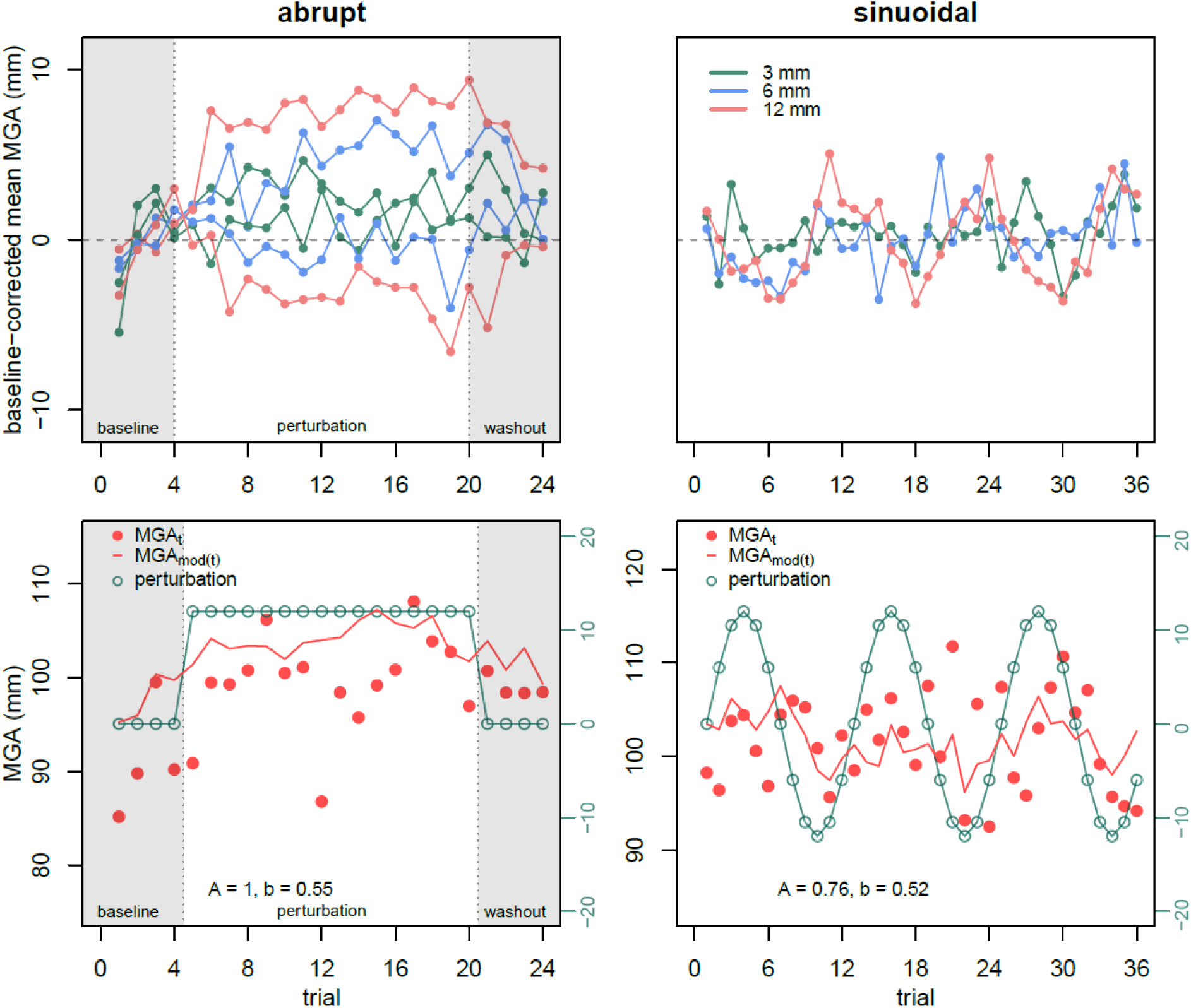
Maximum grip apertures (MGAs) in different conditions. Top: Mean MGAs, split up by perturbation type (abrupt on the left, sinusoidal on the right) and perturbation magnitude (by color), baseline-corrected relative to non-perturbed trials. Bottom: MGAs, perturbations and the corresponding models from sample blocks. Models computed following equation 1. We show data from individual blocks rather than aggregates since the use of the trial-wise MGA, along with a block-wise correction parameter, makes an average model fitted to average data hard to interpret. Left: Data from block with an abrupt perturbation. Right: Data from a sinusoidal-perturbation block. For the top row, absolute values relative to baseline were computed to enable collapsing of blocks using sinus function with different signs.

### Discrimination performance

Participants were slightly more likely to judge the haptic object as larger than the visual one (56.8%), and also more likely to respond that the object was definitely larger or smaller (64.6%) than it being only probably so. Both confidence and size judgements were more sensitive to differences in perturbation magnitude in sinusoidal-perturbation blocks compared to abrupt-perturbation blocks, see Fig. 4.

**Figure 4:**
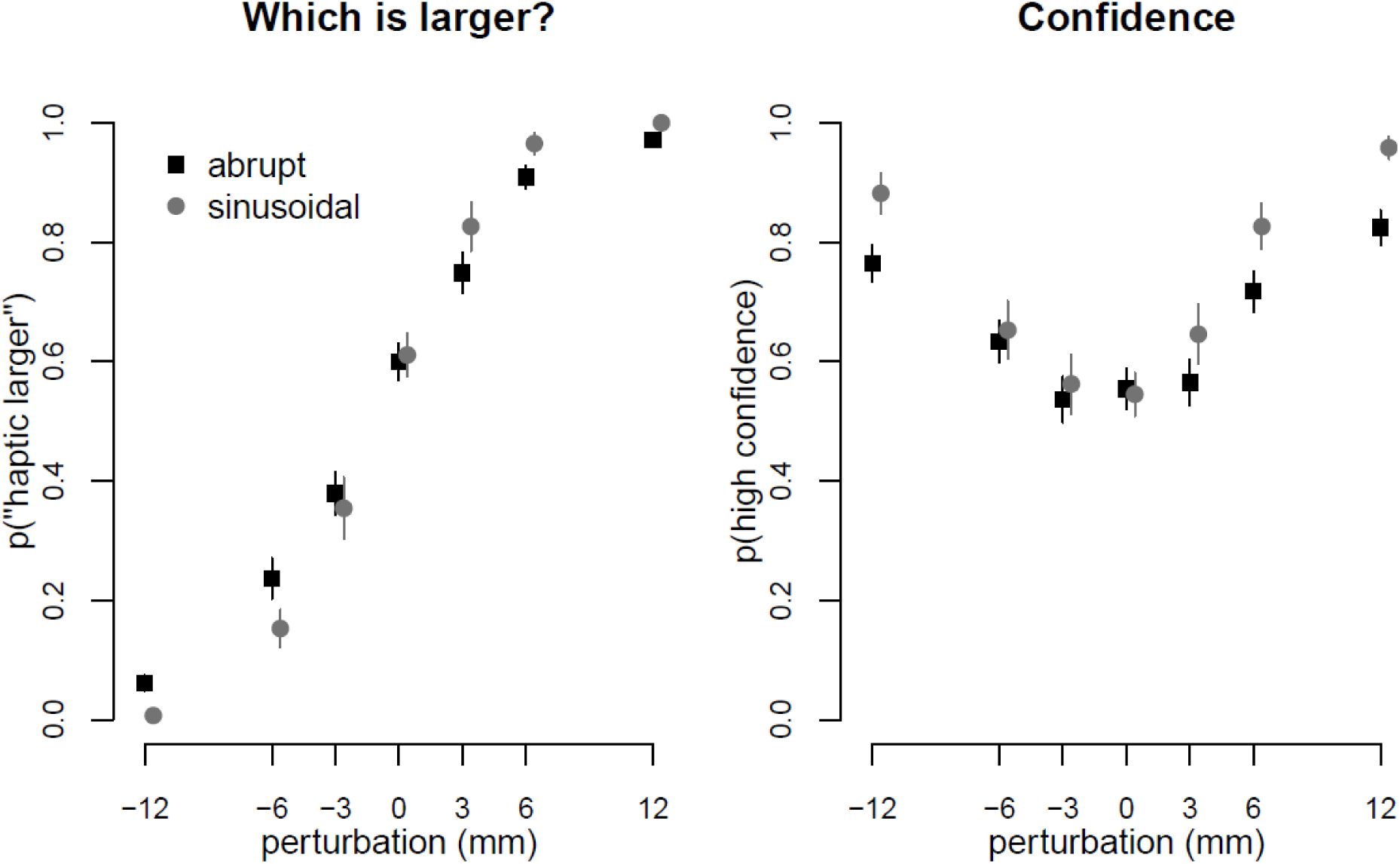
Proportion of responses along the two response dimensions, by perturbation type and magnitude. Left: Proportion of responses saying that the haptic object was larger than the visual one. Right: Proportion of responses with high confidence. Black squares indicate abrupt-perturbation blocks, gray circles show values from sinusoidal-perturbation blocks (data from maximal-perturbation trials only). We show arithmetic means across participants, with between-participant standard errors as error bars.

We calculated linear slopes for each experimental block for the responses’ correctness (with respect to the direction of the perturbation and binarized, so the combined proportion of dark and light blue, Fig. 5), so *correctness* ∼ *trial number*. These were submitted to a 2x3 rmANOVA, revealing a significant effect for the perturbation condition (sinusoidal/abrupt) on response accuracy: *F*(1, 47) = 48.86, *p* < .001 for the average slope across trials (decreasing by −0.95% per trial for abrupt conditions, and decreasing by −0.02% per trial for sinusoidal conditions), consistent with the fact that visually, the proportion of correct trials in Fig. 5 decreased over trials for abrupt, but not sinusoidal perturbations, as one would expect if sensorimotor adaptation makes detection harder. However, no significant main effect was found for perturbation size (*F*(2, 94) = 2.72, *p* = .071), nor an interaction effect (*F*(2, 94) = 1.88, *p* = .158).

**Figure 5:**
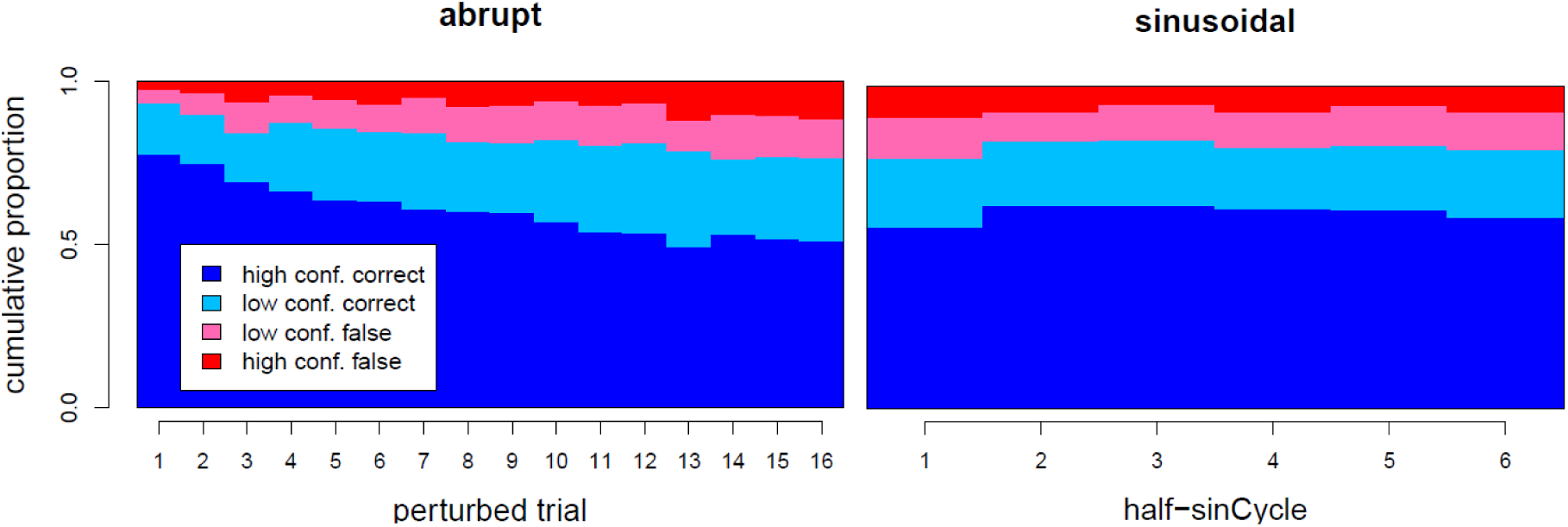
Participants’ responses across trials. The y-axis shows the proportions of each response, cumulatively, such that the height of the dark-blue bar is the proportion of responses that were correct and high-confidence, the height of the light-blue bar on top of this is the proportion of low-confidence responses that were correct, and light red and dark red show the proportions of low-confidence and high-confidence responses that were incorrect. The x-axis displays trials (left) or sinus half-cycles (right). Left: Data from blocks with abruptly-introduced perturbations, with mean proportions plotted by trial, starting with the first perturbed trial. Right: Data from sinusoidal blocks, plotted by sinus-half-cycle, which would contain each perturbation magnitude in the block precisely once and is thus free of the confounding factor of perturbation magnitude.

For linear slopes of response confidence (combined proportion of dark red and dark blue, Fig. 5), the 2x3 rmANOVA showed a significant effect of the perturbation condition (sinusoidal/abrupt): *F*(1, 47) = 53.07, *p* < .001 for the average slope across trials (decreasing by −1.21% per trial for abrupt conditions, and decreasing by −0.05% per trial for sinusoidal conditions). Additionally, significant effects were observed of perturbation size (*F*(2, 94) = 7.98, *p* = .001), and an interaction effect (*F*(2, 94) = 6.90, *p* = .002). The significant interaction suggests that the effect of perturbation size on the slope of response confidence depends on whether the perturbation is abrupt or sinusoidal. Similar to the effects seen in the correctness of responses, the proportion of high-confidence responses (and especially high-confidence correct responses) decreased substantially during abrupt, but not sinusoidal-perturbation blocks (Fig. 5).

To investigate and compare overall (aggregated) discrimination performance, we constructed ROC curves for each participant and each perturbation type (Fig. 6) and analyzed the area under the curve (AUC). On average, participants did quite well in the task, with mean AUCs of .88 for abrupt blocks and .93 for sinusoidal blocks. This difference was statistically significant, *t*(47) = 4.8, *p* < .001. We also computed just-noticeable differences from binary-coded responses, finding JND values of 5.43 ± 0.23 mm for the abrupt condition and 3.97 ± 0.17 mm for the sinusoidal condition.

**Figure 6:**
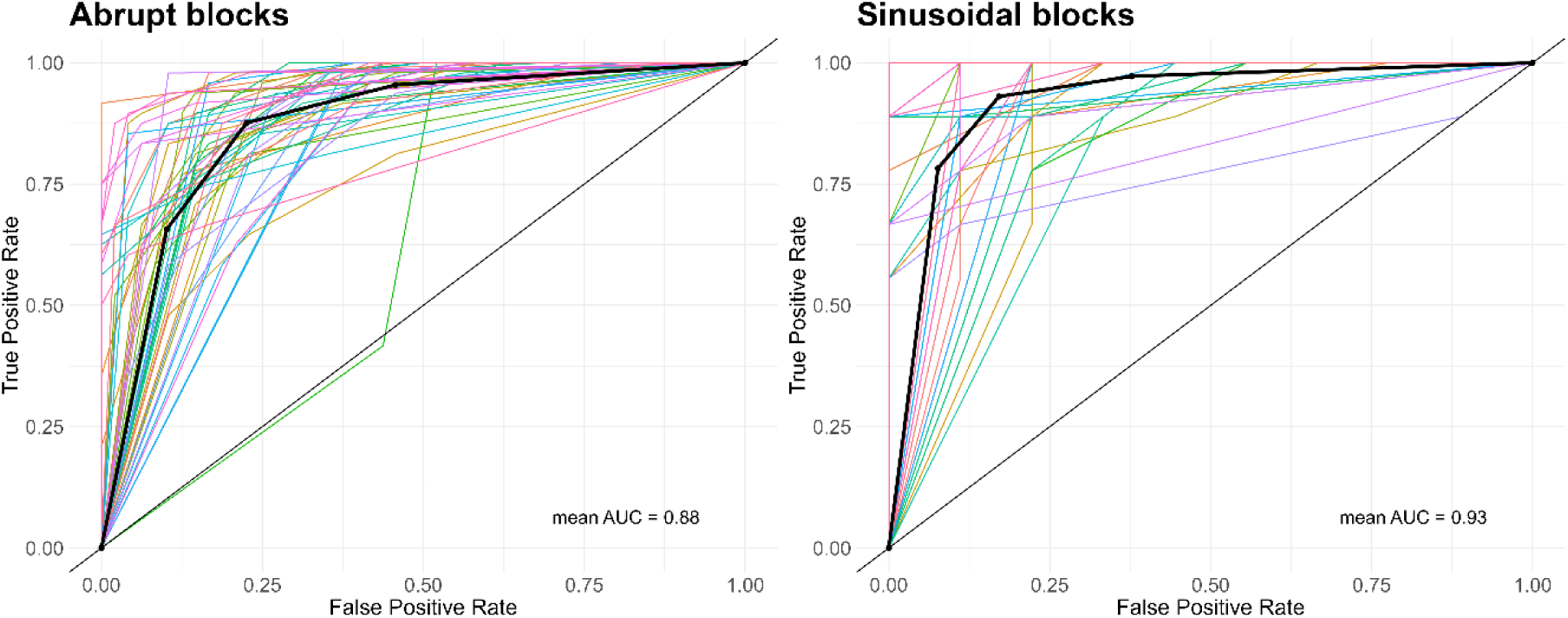
ROC curves for the two types of blocks. Left: Data from abrupt-perturbation blocks, right: Data from sinusoidal-perturbation blocks, maximum-amplitude trials only. Each colored line indicates a single participant, thick black lines indicate ROC curves computed from grand means across participants.

### Pupillary responses

Average pupil dilation depending on trial and response characteristics are shown in Fig. 7. From these, pupil parameters baseline, velocity difference, and amplitude were computed for each trial.

**Figure 7:**
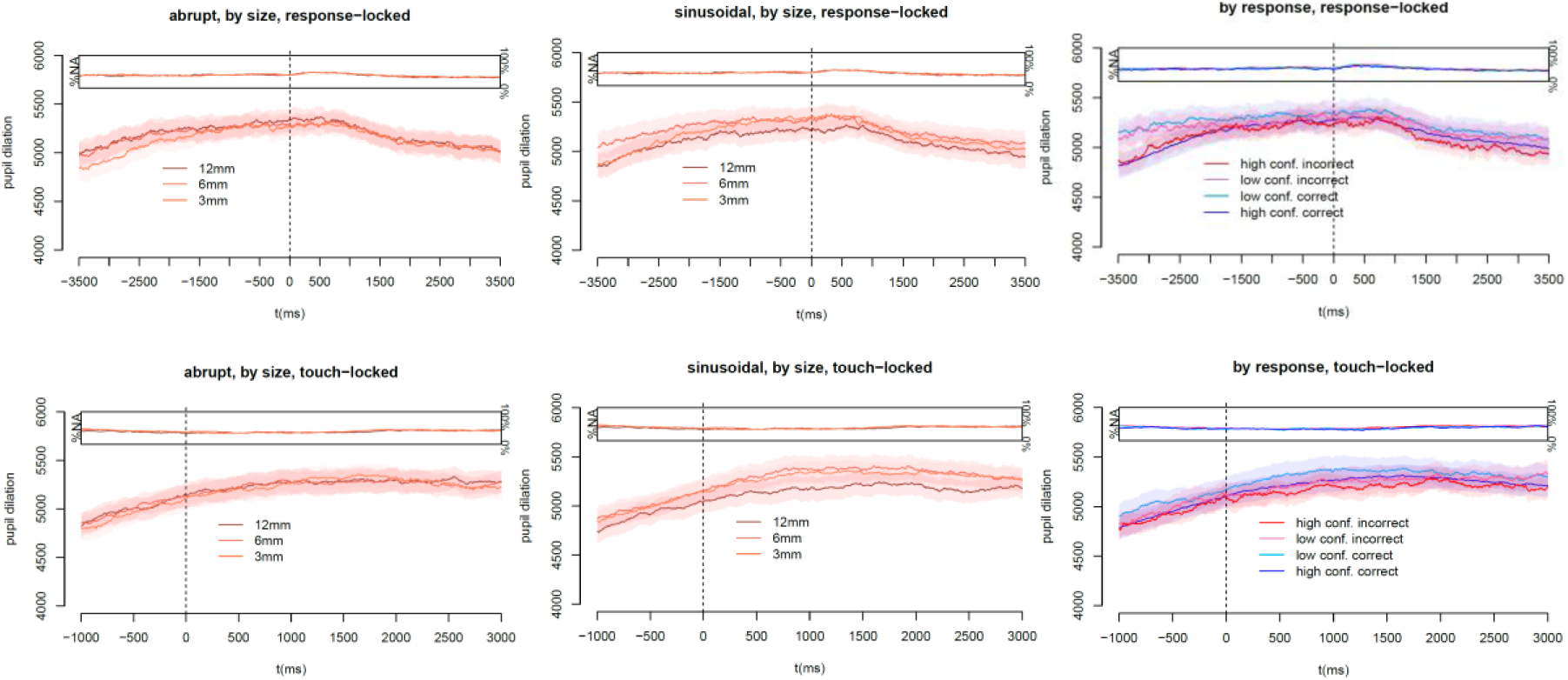
Pupil-dilation trajectories split up by perturbation type and by response. Plotted are grand means across participants. Top row: Trajectories relative to the participant’s response. Bottom row: Trajectories relative to the participant touching the haptic object. Shaded areas indicate ± one between-participant SEM. Insets show proportion of frames with missing data.

Using LMEs, we investigated if these parameters varied systematically depending on (i) perturbation magnitude, (ii) perturbation type, (iii) trial, (iv) block, (v) response correctness, (vi) response confidence, and (vii) the grasping error E. Each of these fixed effects were fit as slopes in the LMEs. Iteratively adding predictors and comparing them by AIC to the previously best-fitting model revealed the best model to be the full model for pupil-dilation *baseline* and for the dilation’s *amplitude*, though we note that some predictors did not affect both variables in the same direction, i.e., one positively and one negatively. For its *velocity difference*, neither trial number nor response confidence improved the model fit (Table 1). Thus, pupil parameters responded to stimulus differences and differed by response and grasping parameters, although with slight differences between tonic and phasic parameters.

**Table 1:**
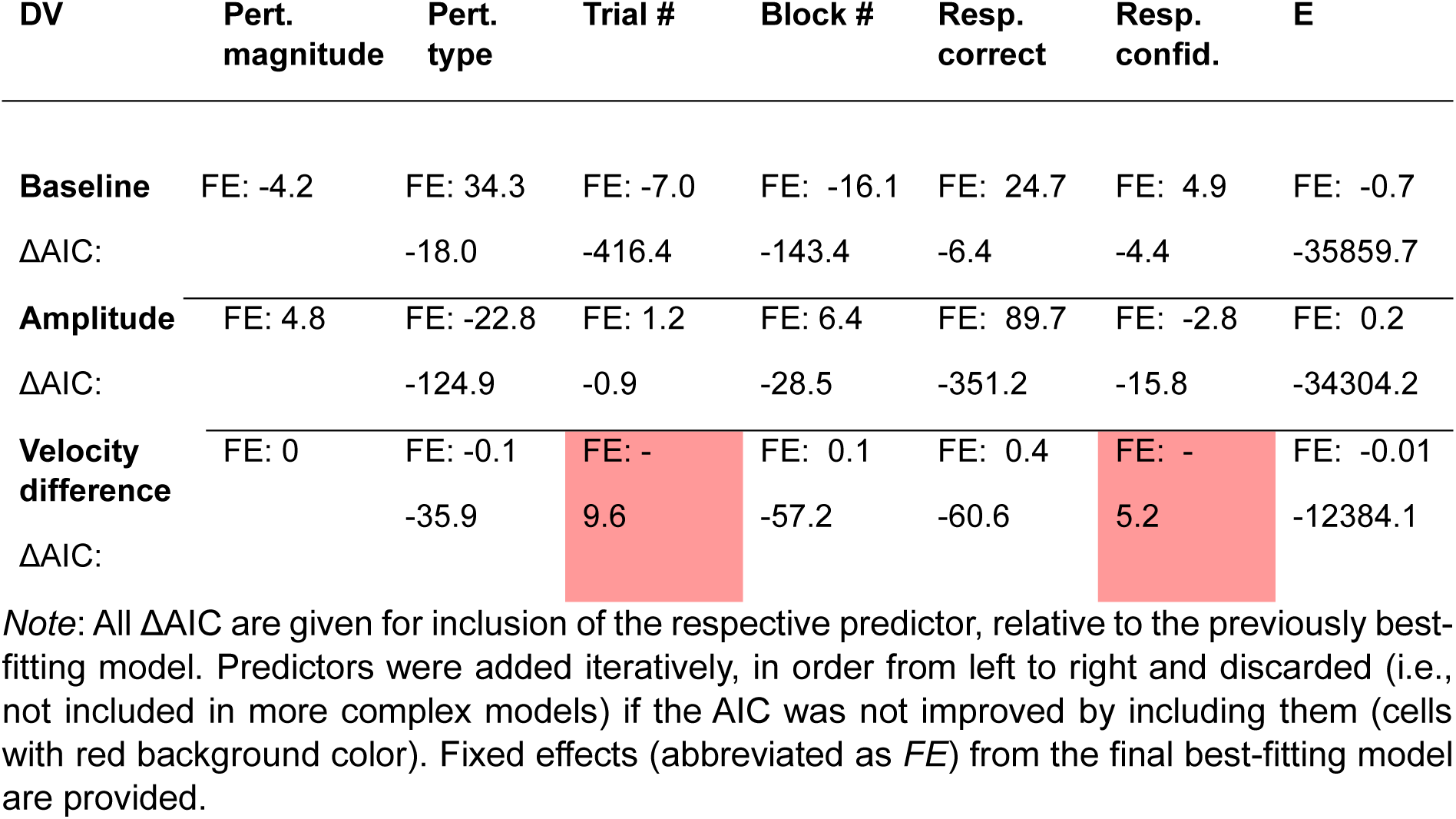
Eye parameters predicted by trial-information variables.

### Classification of response confidence with SVM

Classification performance using our standard set of meta-parameters (50 Hz, C = 1, equal class weights) is summarized in Table 2. As we can see, while class-based accuracy in the training set was quite good for many different sets of features and especially for those involving the derivative of pupil dilation, only classifiers using trial-information features could predict responses in the test set. Indeed, adding other features like pupil dilation, its derivative, or grasping error to a model containing trial information reliably increased the accuracy in the training set, but *decreased* accuracy in the test set.

**Table 2:** SVM results with pupil dilation and its derivative sampled at 50 Hz.

| Model | Acc. Train | Acc. Test | nSamples Train | nSamples Test |
| --- | --- | --- | --- | --- |
| **Trial information** | 90.5% | 74.5% | 105.1 | 32.1 |
| **Eye parameters** | 67.5% | 47.4% | 94.5 | 32.1 |
| **Dilation trajectory** | 72.4% | 48.7% | 100.1 | 27.7 |
| **Dilation derivative** | 96.2% | 47.4% | 99.1 | 28.8 |
| **Dilation trajectory + derivative** | 90.5% | 49.0% | 95.6 | 27.8 |
| **Dilation trajectory + derivative + info** | 94.8% | 50.0% | 100.3 | 25.7 |
| **Grasping error** | 63.2% | 50.4% | 99.2 | 31.0 |
| **Grasping error + trial info** | 91.6% | 71.9% | 99.2 | 29.3 |

*Note:* We report arithmetic means across participants. “Accuracy” refers to class-balanced accuracy. The number of samples differed as trials with missing values in the features were excluded.

The obvious possible explanation here is overfitting: With 50 Hz, we had several times as many features if we used the dilation trajectories than we had trials in the training set. Thus, conducted the same analysis with lower sampling rates of 10 Hz and 5 Hz, see Table A.1 and Table A.2. The main difference here was that while the overall pattern stayed the same – only trial-information features having any predictive power in the test set – the accuracy in the test set tended to decrease less than with the higher sampling rate. Thus, it is likely that (i) overfitting was indeed a problem, and (ii) the information contained in most features was not sufficient for classification. A multiverse analysis varying not only framerate but also the class weights and cost factor (Table A.3) showed the same pattern, with only models including trial information performing consistently above chance in the test set, and none outperforming the simple trial-information-only model.

## Discussion

We found that as participants adapted to a sensorimotor size perturbation in grasping, their discrimination performance regarding the same perturbation magnitude decreased, as did the confidence in their own responses. We replicated and extended previous work (Müller et al., 2025), who also found reduced discrimination with abrupt perturbations, and generalized the results to a four-response setting (thereby circumventing methodological problems of participants repeatedly having to give the same response). We also probed whether pupil responses could be used to predict the confidence of participants’ responses, as a first step towards using pupillometry as a no-report marker of perturbation detection. While we did find that pupil parameters responded to not only differences between experimental trials (Yokoi & Weiler, 2022), but also differed depending on participants’ psychophysical and grasping responses, using pupil information as features in an SVM classifier did not allow us to accurately predict psychophysical responses.

Having previously found that perturbation schedules that are easy to adapt to correlate with decreasing perturbation detection (Müller et al., 2025) in a 2AFC task, part of our study was aimed at improving the prior study methodologically. A major concern about presenting an abruptly introduced step-function perturbation is that it requires participants to give the same answer many times in a row, potentially introducing response biases that cannot be dissociated from the putative effects of sensorimotor adaptation on detection. This was ameliorated by our use of a four-response task, as participants had multiple correct options on any given perturbation trial. Not only did we replicate the performance decrease with respect to correctness, but we also found the same pattern in response confidence, again in line with the idea that it is sensorimotor adaptation itself that, by decreasing the grasping error, makes it harder to detect the perturbation participants adapted to and participants more uncertain about their responses.

Our experiments focused on responses – motor, psychophysical, and physiological – to motor perturbations, that is, externally induced errors. This is distinct from responses to self-generated errors, which may be the more common type of error in everyday life. Here, recent work on metacognition has shown that participants are also able to judge errors that are not induced by the experimenter (Arbuzova et al., 2021) at an above-chance rate. Interestingly, such metacognition appears to be preserved in confidence ratings even when detection responses are incorrect (Pereira et al., 2023). While we found no difference in patterns between the correctness and confidence of responses (both affected similarly on average by the perturbations and decreasing over time for abrupt but not sinusoidal perturbation schedules), experiments targeted at investigating metacognition over time may be an interesting avenue to find out more about what is used to make metacognitive judgements. More generally, the question is to what extent the results here generalize to other settings, which includes other motor actions such as reaching (Gaffin-Cahn et al., 2019) or walking (Iturralde et al., 2020; Müller & Kopiske, 2025), as well as psychophysical tasks more specifically designed to assess participants’ confidence (Fleming & Lau, 2014; Mamassian, 2016).

Finally, we show that pupil dilation reflects not just the characteristics of experimental trials (Yokoi & Weiler, 2022), but also characteristics of participants’ responses. LME analyses confirmed that both the tonic response, quantified here by baseline dilation before the start of each trial, and the phasic response, quantified as dilation change after touching the haptic object, depended on trial characteristics such as the perturbation as well as trial and block number, and on response characteristics such as the correctness and confidence of responses in the detection task and the grasping error. In line with previous results (Yokoi & Weiler, 2022), this is consistent with involvement of noradrenaline and the locus coeruleus (Dayan & Yu, 2006) as the actor acts and decides under uncertainty. Such effects are a necessary condition for the overarching long-term goal: Predicting psychophysical responses from pupil data. In simple terms, for this to be possible, pupil dilation and psychophysics need to be related at all, which is what the LMEs demonstrate. To go one step further, we also trained SVM classifiers using different sets of features – trial information, aggregated pupil-dilation parameters, pupil-dilation trajectories, and grasping errors – to predict response confidence. This is in line with previous work arguing that pupil responses can reflect uncertainty and conflicting information (Ebitz & Platt, 2015; Joshi & Gold, 2020), which is why we attempted to predict response confidence rather than correctness (which participants also did not have access to when they gave their responses).

This was only partially successful: While many models showed great accuracy (>85%) in the training set, only trial-information features had any predictive power in the test set, and indeed, models containing these features only performed substantially better on the test set than those additionally containing other features. In particular, the derivative of pupil-dilation trajectories performed exceptionally well in the training set, but at chance level in the training set, even combined with trial-information features. Using fewer features by downsampling pupil trajectories to combat overfitting ameliorated the latter problem, but still the pupil data showed no benefit over just using trial information. Here, we note three things: One, it is possible that improved data quality could improve classification, although the mean proportion of missing data was only moderately worse in trials that were incorrectly classified (44.6%) compared to those that were correctly classified (41.2%) using pupil-dilation features. Two, while we chose to temporally lock trajectories to the participant touching the object, the time courses plotted in Fig. 7 suggest that locking them to the participant response might be just as promising – however, investigating the time course in detail warrants its own study and is beyond the scope of this manuscript. Three, we deliberately used SVM as a standard, well-tested classifier. Our study focused on whether there was something in the data, not how well cutting-edge machine-learning classifiers can perform. In future research, it may be useful to take a step back and verify if, in a simple adaptation paradigm without psychophysical response, trial characteristics can be predicted from pupil data.

## Conclusion

As humans adapt to motor perturbations, the motor error decreases. This, in turn, makes it harder for them to detect those same perturbations, and makes them less confident in being able to do so. Pupil-dilation parameters responded to trial- and response-characteristics, but did not allow accurate classification of participant responses.

## Acknowledgements

These data were also presented at the 2024 European Conference on Visual Perception. We thank Wolfgang Einhäuser-Treyer for his helpful comments on the manuscript. This work was supported by a grant from the German Research Foundation (DFG) to KK (DFG KO 6478-1/1; project number 466287772).

## Appendix

**Table A.1:** SVM results with pupil dilation and its derivative sampled at 10 Hz.

| **Model** | **Acc. Train** | **Acc. Test** | **nSamples Train** | **nSamples Test** |
| --- | --- | --- | --- | --- |
| **Trial information** | 90.0% | 78.8% | 105.0 | 33.8 |
| **Eye parameters** | 67.5% | 45.4% | 96.4 | 28.2 |
| **Dilation trajectory** | 72.6% | 50.1% | 96.8 | 29.7 |
| **Dilation derivative** | 92.9% | 48.4% | 98.3 | 30.1 |
| **Dilation trajectory + derivative** | 88.7% | 45.5% | 101.3 | 28.0 |
| **Dilation trajectory + derivative + info** | 96.0% | 62.6% | 96.3 | 28.6 |
| **Grasping error** | 65.6% | 46.9% | 96.6 | 30.2 |
| **Grasping error + trial info** | 91.0% | 73.9% | 102.7 | 28.3 |

*Note:* Variables used as in Table 2.

**Table A.2:** SVM at 5 Hz.

| **Model** | **Acc. Train** | **Acc. Test** | **nSamples Train** | **nSamples Test** |
| --- | --- | --- | --- | --- |
| **Trial information** | 91.4% | 74.4% | 105.4 | 29.5 |
| **Eye parameters** | 67.9% | 41.0% | 99.2 | 28.2 |
| **Dilation trajectory** | 72.1% | 45.6% | 100.1 | 27.1 |
| **Dilation derivative** | 89.4% | 49.6% | 97.0 | 27.0 |
| **Dilation trajectory + derivative** | 87.6% | 48.5% | 97.6 | 28.3 |
| **Dilation trajectory + derivative + info** | 96.1% | 69.6% | 98.6 | 27.6 |
| **Grasping error** | 65.6% | 54.4% | 99.0 | 29.8 |
| **Grasping error + trial info** | 91.8% | 73.3% | 102.4 | 28.1 |

*Note:* Variables used as in Table 2.

**Table A.3:** Multiverse SVM results.

| Model | Accuracy Training | Accuracy Test |
| --- | --- | --- |
| **Trial information** | 75.3% - 98.5% | 55.8% - 100% |
| **Eye parameters** | 59.8% - 82.1% | 24.7% - 52.8% |
| **Dilation trajectory** | 59.2% - 100% | 39.0% - 75.9% |
| **Dilation derivative** | 75.1% - 99.8% | 37.5% - 63.0% |
| **Dilation trajectory + derivative** | 72.2% - 100% | 39.6% - 64.1% |
| **Dilation trajectory + derivative + info** | 73.3% - 100% | 34.9% - 69.6% |
| **Grasping error** | 56.9% - 72.6% | 46.6% - 67.3% |
| **Grasping error + trial info** | 73.2% - 100% | 48.8% - 100% |

*Note:* Variables used as in Table 2. We report ranges of means across parameter combinations. Frame rate (50 Hz, 10 Hz, 5 Hz), class weights (equal, inverse), and cost factor ( $C = 10^{-5}$ , $C = 10^{-1}$ , $C = 1$ , $C = 10$ , $C = 10^5$ ) were varied independently.

## References

Akaike, H. (1974). A new look at the statistical model identification. IEEE Transactions on Automatic Control AC*-*19, 716–723.

Arbuzova, P., Peters, C., Röd, L., Koß, C., Maurer, L. K., Müller, H., Verrel, J., & Filevich, E. (2021). Measuring metacognition of direct and indirect parameters of voluntary movement. Journal of Experimental Psychology: General, 150(11), 2208–2219. 10.1037/xge0000892

Bates, D., Mächler, M., Bolker, B., & Walker, S. (2015). Fitting linear mixed-effects models using lme4. Journal of Statistical Software, 67(1), 1–51. 10.18637/jss.v067.i01

Bhatia, K., Löwenkamp, C., & Franz, V. H. (2022). Grasping follows Weber’s law: How to use response variability as a proxy for JND. Journal of Vision, 22(12), 13. 10.1167/jov.22.12.13

Bosch, E., Fritsche, M., Ehinger, B. V., & de Lange, F. P. (2020). Opposite effects of choice history and evidence history resolve a paradox of sequential choice bias. Journal of Vision, 20(*12*)(9), 1–13. 10.1167/jov.20.12.9

Boser, B. E., Guyon, I. M., & Vapnik, V. N. (1992). A training algorithm for optimal margin classifiers. Proceedings of the Fifth Annual Workshop on Computational Learning Theory, 144–152. 10.1145/130385.130401

Burnham, K. P., & Anderson, R. P. (2004). Multimodel Inference: Understanding AIC and BIC in Model Selection. Sociological Methods & Research, 33(2), 261–304. 10.1177/0049124104268644

Cesanek, E., & Domini, F. (2017). Error correction and spatial generalization in human grasp control. Neuropsychologia, 106, 112–122. 10.1016/j.neuropsychologia.2017.09.026

Cheng, S., & Sabes, P. N. (2006). Modeling sensorimotor learning with linear dynamical systems. Neuronal Computation, 18(4), 760–793. 10.1162/089976606775774651.Modeling

Cohen, J. (1988). Statistical power analysis for the behavioral sciences (2nd ed.). Psychology Press.

Dayan, P., & Yu, A. J. (2006). Phasic norepinephrine: A neural interrupt signal for unexpected events. Network: Computation in Neural Systems, 17(4), 335–350. 10.1080/09548980601004024

Ebitz, R. B., & Platt, M. L. (2015). Neuronal activity in primate dorsal anterior cingulate cortex signals task conflict and predicts adjustments in pupil-linked arousal. Neuron, 85(3), 628–640. 10.1016/j.neuron.2014.12.053

Einhäuser, W., Stout, J., Koch, C., & Carter, O. (2008). Pupil dilation reflects perceptual selection and predicts subsequent stability in perceptual rivalry. Proceedings of the National Academy of Sciences, 105(5), 1704–1709. 10.1073/pnas.0707727105

Fleming, S. M., & Lau, H. C. (2014). How to measure metacognition. Frontiers in Human Neuroscience, 8. 10.3389/fnhum.2014.00443

Gaffin-Cahn, E., Hudson, T. E., & Landy, M. S. (2019). Did I do that? Detecting a perturbation to visual feedback in a reaching task. Journal of Vision, 19(1)(5), 1–18. 10.1167/19.1.5

Gentilucci, M., Daprati, E., Toni, I., Chieffi, S., & Saetti, M. C. (1995). Unconscious updating of grasp motor control. Experimental Brain Research, 105, 291–303.

Green, D. M., & Swets, J. A. (1966). Signal detection theory and psychophysics (1st ed.). John Wiley & Sons.

Hewitson, C. L., Al-Fawakhiri, N., Forrence, A. D., & McDougle, S. D. (2023). Metacognitive judgments during visuomotor learning reflect the integration of error history. Journal of Neurophysiology, 130(2), 264–277. 10.1152/jn.00022.2023

Hudson, T. E., & Landy, M. S. (2012). Measuring adaptation with a sinusoidal perturbation function. Journal of Neuroscience Methods, 208, 48–58.

Iturralde, P. A., Gonzalez-Rubio, M., & Torres-Oviedo, G. (2020). *High-human acuity of speed asymmetry during walking* [Preprint]. Bioengineering. 10.1101/2020.10.28.359281

Jeannerod, M. (1984). The timing of natural prehension movements. Journal of Motor Behavior, 16(3), 235–254.

Joshi, S., & Gold, J. I. (2020). Pupil size as a window on neural substrates of cognition. Trends in Cognitive Sciences, 24(6), 466–480. 10.1016/j.tics.2020.03.005

Kopiske, K. K., Cesanek, E., Campagnoli, C., & Domini, F. (2017). Adaptation effects in grasping the Müller-Lyer illusion. Vision Research, 136, 21–31. 10.1016/j.visres.2017.05.004

Krakauer, J. W., & Mazzoni, P. (2011). Human sensorimotor learning: Adaptation, skill, and beyond. Current Opinion in Neurobiology, 21(4), 636–644. 10.1016/j.conb.2011.06.012

Kuhn, M., & Johnson, K. (2013). Applied Predictive Modeling. Springer New York. 10.1007/978-1-4614-6849-3

Mamassian, P. (2016). Visual confidence. Annual Review of Vision Science, 2(1), 459–481. 10.1146/annurev-vision-111815-114630

Mariscal, D. M., Iturralde, P. A., & Torres-Oviedo, G. (2020). Altering attention to split-belt walking increases the generalization of motor memories across walking contexts. Journal of Neurophysiology, 123, 1838–1848. 10.1152/jn.00509.2019

Mazzoni, P., & Krakauer, J. W. (2006). An implicit plan overrides an explicit strategy during visuomotor adaptation. Journal of Neuroscience, 26(14), 3642–3645. 10.1523/JNEUROSCI.5317-05.2006

McDougle, S. D., Bond, K. M., & Taylor, J. A. (2015). Explicit and implicit processes constitute the fast and slow processes of sensorimotor learning. Journal of Neuroscience, 35(26), 9568– 9579. 10.1523/JNEUROSCI.5061-14.2015

McDougle, S. D., Ivry, R. B., & Taylor, J. A. (2016). Taking aim at the cognitive side of learning in sensorimotor adaptation tasks. Trends in Cognitive Sciences, 20(7), 535–544. 10.1016/j.tics.2016.05.002

McDougle, S. D., Wilterson, S. A., Turk-Browne, N. B., & Taylor, J. A. (2022). Revisiting the role of the medial temporal lobe in motor learning. Journal of Cognitive Neuroscience, 34(3), 532–549. 10.1162/jocn_a_01809

Meyer, D., Dimitriadou, E., Hornik, K., Weingessel, A., Leisch, F., Chang, C. C., & Lin, C. C. (2024). *E1071* (Version 1.7-16) [Computer software]. https://doi.org10.32614/CRAN.package.e1071

Miall, R. C., & Wolpert, D. M. (1996). Forward models for physiological motor control. Neural Networks, 9(8), 1265–1279. 10.1016/S0893-6080(96)00035-4

Milgram, P. (1987). A spectacle-mounted liquid-crystal tachistoscope. *Behavior Research Methods*, Instruments & Computers, 19(5), 449–456.

Miyamoto, Y. R., Wang, S., & Smith, M. A. (2020). Implicit adaptation compensates for erratic explicit strategy in human motor learning. Nature Neuroscience, 23, 443–455. 10.1038/s41593-020-0600-3

Modchalingam, S., Ciccone, M., D’Amario, S., ’t Hart, B. M., & Henriques, D. Y. P. (2023). Adapting to visuomotor rotations in stepped increments increases implicit motor learning. Scientific Reports, 13(1), 5022. 10.1038/s41598-023-32068-8

Müller, C., Bendixen, A., & Kopiske, K. (2025). Sensorimotor adaptation impedes perturbation detection in grasping. Psychonomic Bulletin & Review, 32, 373–386. 10.3758/s13423-024-02543-y

Müller, C., & Kopiske, K. (2025). Perceiving inter-leg speed differences while walking on a split-belt treadmill. Scientific Reports, 15(1), 1375. 10.1038/s41598-024-85091-8

Naber, M., Frässle, S., & Einhäuser, W. (2011). Perceptual rivalry: Reflexes reveal the gradual nature of visual awareness. PLoS ONE, 6(6), e20910. 10.1371/journal.pone.0020910

Naber, M., Frässle, S., Rutishauser, U., & Einhäuser, W. (2013). Pupil size signals novelty and predicts later retrieval success for declarative memories of natural scenes. Journal of Vision, 13(2), 11–11. 10.1167/13.2.11

Orban de Xivry, J. J., Ahmadi-Pajouh, M. A., Harran, M. D., Salimpour, Y., & Shadmehr, R. (2013). Changes in corticospinal excitability during reach adaptation in force fields. Journal of Neurophysiology, 109, 124–136. 10.1152/jn.00785.2012

Pereira, M., Skiba, R., Cojan, Y., Vuilleumier, P., & Bègue, I. (2023). Preserved metacognition for undetected visuomotor deviations. The Journal of Neuroscience, 43(35), 6176–6184. 10.1523/JNEUROSCI.0133-23.2023

Säfström, D., & Edin, B. B. (2005). Short-term plasticity of the visuomotor map during grasping movements in humans. Learning & Memory, 12(1), 67–74. 10.1101/lm.83005

Savitzky, A., & Golay, M. J. E. (1964). Smoothing and differentiation of data by simplified least squares procedures. Analytical Chemistry, 36(8), 1627–1639. 10.1021/ac60214a047

Shadmehr, R., Smith, M. A., & Krakauer, J. W. (2010). Error correction, sensory prediction, and adaptation in motor control. Annual Review of Neuroscience, 33, 89–108.

Smeets, J. B. J., & Brenner, E. (1999). A new view on grasping. Motor Control, 3(3), 237–271.

Taylor, J. A., & Ivry, R. B. (2011). Flexible cognitive strategies during motor learning. PLoS Computational Biology, 7(3). 10.1371/journal.pcbi.1001096

Taylor, J. A., Krakauer, J. W., & Ivry, R. B. (2014). Explicit and implicit contributions to learning in a sensorimotor adaptation task. Journal of Neuroscience, 34(8), 3023–3032. 10.1523/JNEUROSCI.3619-13.2014

Warren, W. H. (2006). The dynamics of perception and action. Psychological Review, 113(2), 358–389. 10.1037/0033-295X.113.2.358

Wolpert, D. M., Ghahramani, Z., & Jordan, M. I. (1995). An internal model for sensorimotor integration. Science, 269, 1880–1882. 10.1126/science.7569931

Yokoi, A., & Weiler, J. (2022). Pupil diameter tracked during motor adaptation in humans. Journal of Neurophysiology, 128(5), 1224–1243. 10.1152/jn.00021.2022

Ypma, J. (2014). nloptr: R interface to NLopt. https://cran.r-project.org/package=nloptr

